# Consensus tissue domain detection in spatial multi-omics data using MILWRM

**DOI:** 10.1101/2023.02.02.526900

**Authors:** Harsimran Kaur, Cody N. Heiser, Eliot T. McKinley, Lissa Ventura-Antunes, Coleman R. Harris, Joseph T. Roland, Martha J. Shrubsole, Robert J. Coffey, Ken S. Lau, Simon Vandekar

## Abstract

Spatially resolved molecular assays provide high dimensional genetic, transcriptomic, proteomic, and epigenetic information in situ and at various resolutions. Pairing these data across modalities with histological features enables powerful studies of tissue pathology in the context of an intact microenvironment and tissue structure. Increasing dimensions across molecular analytes and samples require new data science approaches to functionally annotate spatially resolved molecular data. A specific challenge is data-driven cross-sample domain detection that allows for analysis within and between consensus tissue compartments across high volumes of multiplex datasets stemming from tissue atlasing efforts. Here, we present MILWRM – multiplex image labeling with regional morphology – a Python package for rapid, multi-scale tissue domain detection and annotation. We demonstrate MILWRM’s utility in identifying histologically distinct compartments in human colonic polyps and mouse brain slices through spatially-informed clustering in two different spatial data modalities. Additionally, we used tissue domains detected in human colonic polyps to elucidate molecular distinction between polyp subtypes. We also explored the ability of MILWRM to identify anatomical regions of mouse brain and their respective distinct molecular profiles.

## Introduction

The advent of spatially resolved molecular assays has enabled access to high dimensional genetic, transcriptomic, proteomic’ and even epigenetic information in situ while preserving the spatial information lost in single-cell or bulk molecular assays (17, 23, 42, 50). Spatially resolved data can provide powerful insight into interactions between cell types’ progressive changes in tissue architecture in diseases such as cancer, or interactions between different structures in tissue such as lymphoid follicles and blood vessels (29, 48, 53). Biological insights can be derived from recurring spatial patterns extracted using quantitative analysis on spatial data.

Many current methods attempt to complement single-cell analyses, essentially taking a bottom-up approach to reconstruct tissue domains, architectures, and communities from individual cells. In general, individual cells can be identified from high dimensional imaging data by segmentation. Cellular segmentation and annotation are the most challenging step in this kind of approach. There are various methods available for cellular segmentation (25, 40), annotation (39) and neighborhood analysis (20, 34, 59). Widely used lower resolution imaging data such as spatial transcriptomics (ST) and imaging mass spectrometry data are analyzed using cellular deconvolution algorithms to approximate singlecell composition. Most of these algorithms require a parallel single-cell dataset for use as reference (11, 18). Different cell types are then arranged into interaction networks based on their spatial distributions and/or molecular interactions, and these networks are assembled into larger spatial structures that identify tissue-or organ-level domains. This type of analysis has been used for identifying cellular communities in various cancer types associated with patient prognosis (12, 31, 33, 51).

Another perspective comes from the pathology field, where spatial domains and architectures are first identified, followed by instances of cell identification by morphology, which is known as the top-down approach (38). Since this approach focuses directly on pixel-level information instead of reconstruction from single-cell data, it can identify both extracellular structures and cellular communities over a range of micro-to macro-scale. Pixel-based analysis also forms the basis of modern artificial intelligence learning from imaging, and thus paves the way for more complex learning algorithms to be applied to multiplex tissue data (10, 45, 61).

Various methods are currently available for pixel-based spatial domain detection from ST data (8, 55, 63). However, they lack the scalability to work across batches and samples. Attempts to apply these methods across samples fail to yield global consensus domains, and instead identify regional domains that are sample-specific or confounded by batch effects. To decipher true emergent properties within spatial tissue domains, it is imperative that findings can be generalized across many samples at different resolutions. Here, we present multiplex image labeling with regional morphology (MILWRM) that is designed specifically for consensus tissue domain characterization across large sample sets with potential differing orientations and resolutions.

## Results

### The MILWRM pipeline generates consensus tissue domains across specimens

Whereas most spatial analysis algorithms focus on individual specimens, MILWRM aims to identify consensus tissue domains across samples with spatially resolved molecular data (e.g., multiplexed immunofluorescence [mIF] and ST). The MILWRM pipeline can be broadly categorized into three major steps: data preprocessing, tissue domain identification, and tissue domain analysis (Figure 1). To generalize pixel neighborhood information across batches, data preprocessing incorporates downsampling, normalization, data smoothing, and dimensionality reduction. Preprocessing steps differ slightly for mIF and ST (Methods). After preprocessing, tissue domains are identified using unsupervised K-means clustering by subsampling data across all samples (Methods). The number of tissue domains is adjusted by inertia analysis (21). Each pixel is assigned a tissue domain based on the nearest centroid. Domain profiles are calculated by MILWRM from the initial feature space to molecularly describe each tissue domain, which is useful for downstream annotation. Finally, MILWRM computes a variety of metrics to assess the quality of identified tissue domains (Methods). Overall, MILWRM is a comprehensive, easy to use pipeline for tissue domain detection, providing interpretable results for biological analysis and quality assessment.

**Fig. 1.**
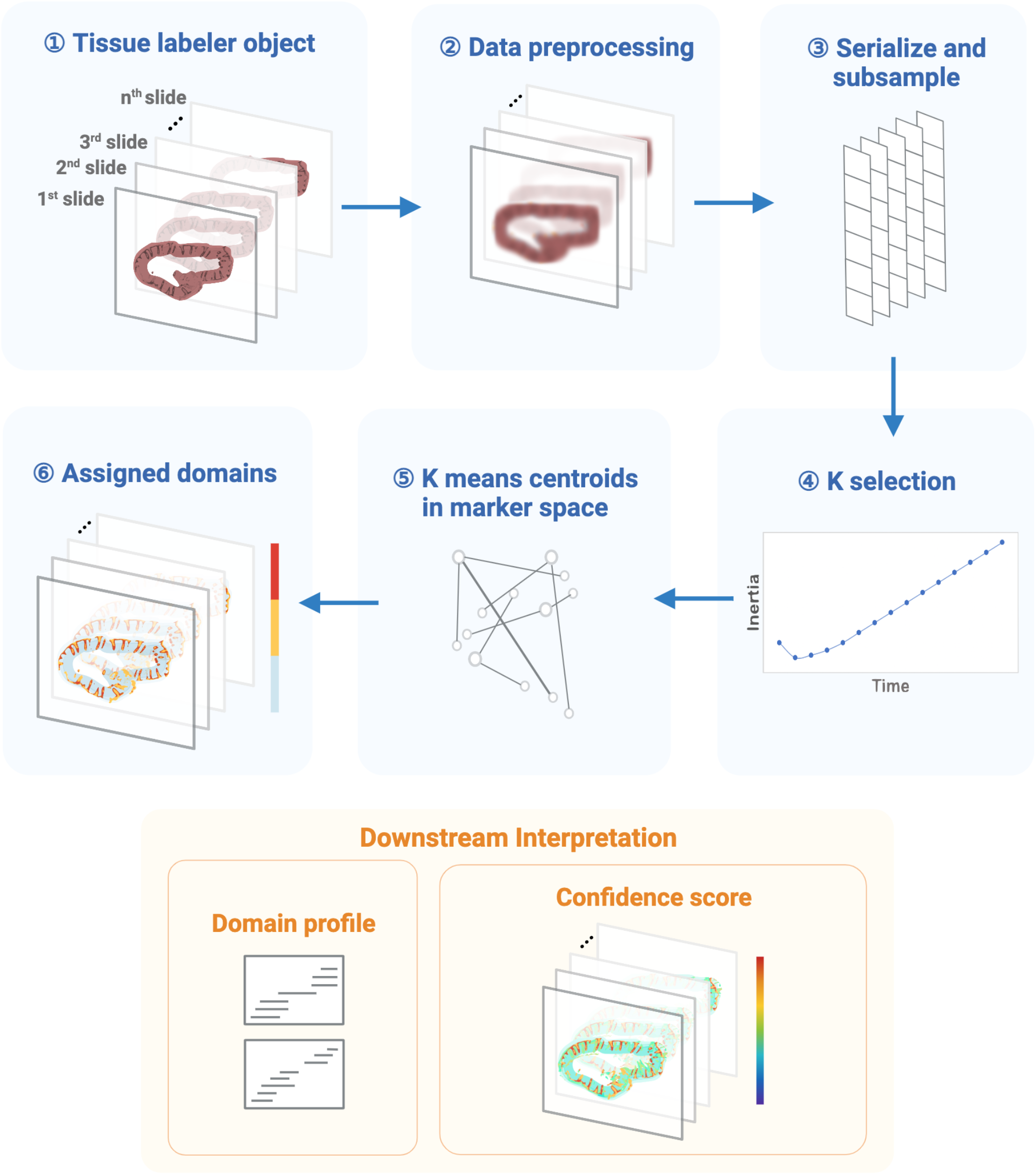
The workflow of MILWRM pipeline. (A) MILWRM begins with constructing a tissue labeler object from all the sample slides that undergoes data preprocessing, serialization and subsampling to create a randomly subsampled dataset used for Kmeans model construction. This subsampled data is used to find optimal number of tissue domains – K selection using adjusted inertia method. Finally, a Kmeans model is constructed, and each pixel is assigned a tissue domain. Each tissue domain has a distinct domain profile describing the molecular feature. MILWRM also provides quality control metrics such as confidence score.

### MILWRM identifies canonical tissue layers of the colonic mucosa

We applied MILWRM to mIF data generated for the Human Tumor Atlas Network (HTAN) consisting of human normal colon and different colonic pre-cancer subtypes (conventional adenomas – AD and serrated polyps – SER) (19). These data comprised multichannel fluorescent images from 37 biospecimens consisting of tissues with different morphologies and pathological classification confirmed by two pathologists (Table S1). We performed low resolution application of MILWRM using a smoothing parameter (*σ*) of 2 after downsampling the images to an isotropic resolution of 5.6 μm/pixel and the penalty parameter of 0.05 that resulted in three tissue domains according to adjusted inertia, as illustrated by three representative samples (Figure 2A-B; Figure S1A). According to domain profiles (Figure 2C), the epithelial monolayer compartment was identified by markers such as CDX2, *β*-catenin, Na+-K+ ATPase, and proliferative marker PCNA, consistent with a high turnover hind-gut epithelium (28, 41). The mucus layer was enriched in MUC2, a secreted mucin (9, 32, 56). The lamina propria region, where stromal cells are prominent, was identified by Vimentin and Collagen (58). The results from MILWRM analysis are consistent with the tissue architecture of the colonic mucosa, as well as other mucosal tissues in the body.

**Fig. 2.**
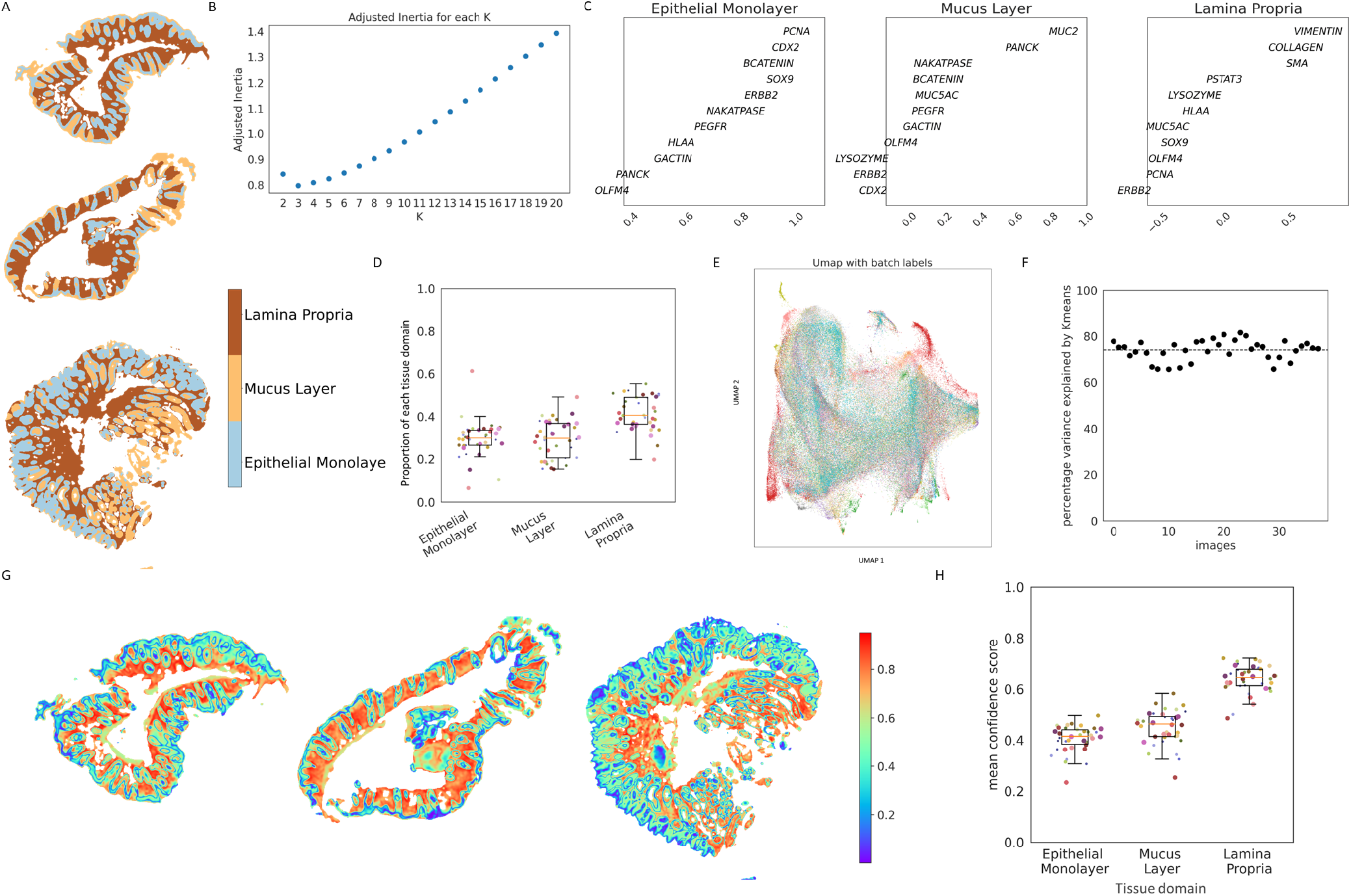
MILWRM detects canonical tissue domains in human colon mIF data. (A) Three representative colon mIF images with labelled tissue domains (*α* = 0.05) (B) Estimated number of tissue domains in Adjusted inertia plot (C) Domain profile describing marker composition of each tissue domain (D) Proportion of each tissue domain across 38 samples (E) UMAP of pixel data used for model building with batch labels (F) Percentage variance explained by Kmeans (G) Three representative colon mIF images with confidence score overlayed (H) mean confidence score in each image for each tissue domain.

MILWRM consistently identified these regions across the 37 tissue samples (Figure 2D), and pixel level data over the samples intermixed in UMAP-embedded space illustrating removal of batch effects between images (Figure 2E), demonstrating the ability of MILWRM to identify consensus regions over multiple samples. MILWRM was able to capture about 80% of the variance in the multidimensional imaging data without any notable outliers, demonstrating that information within the imaging data is retained after MILWRM analysis (Figure 2F). To assess the quality of tissue domain identification, MILWRM calculates a modified silhouette-based confidence score per pixel, which evaluates the deviation of each pixel from the centroid of the matched tissue domain relative to the closest Kmeans centroid. Most pixels across all samples have high confidence scores apart from a few in the epithelial and mucus tissue domains (Figure 2G-H; Figure S2A-B). Low confidence scores can be attributed to inherent biological heterogeneity within epithelial domains, as the analysis is performed over samples from mixed pathological categories (normal, AD, SER). Thus, MILWRM performed on a cohort of 37 biospecimens was able to provide physiologically relevant tissue domains with high confidence.

### MILWRM identifies tissue domains that molecularly distinguish disease subtypes

To obtain more refined tissue domains that appropriately stratify the heterogenous pathological categories of our samples (normal, AD, SER), we next performed MILWRM with a reduced penalty parameter (a = 0.02). We obtained nine tissue domains that further broke down the epithelial compartment into stem (SOX9, PCNA, CDX2), differentiated (Na+-K+ ATPase, PANCK, β-catenin), mucus (MUC2), abnormal (MUC5AC+/PANCK+), and crypt lumen (OLFM4+), and the non-epithelial compartment into smooth muscle, pericryptal stroma, and proximal and deep lamina propria (Figure 3A-C; Figure S3A). These refined tissue domains were spatially localized appropriately. For instance, the stem and crypt lumen regions were located at the crypt base while the differentiated regions were located at the colonic surface. Interestingly, pericryptal stroma was identified with a mixture of epithelial and stromal markers and labeled a thin layer of fibroblasts that comprise telocytes constituting the stem cell niche (Figure 3A-C; Figure S4A-E) (13, 52).

**Fig. 3.**
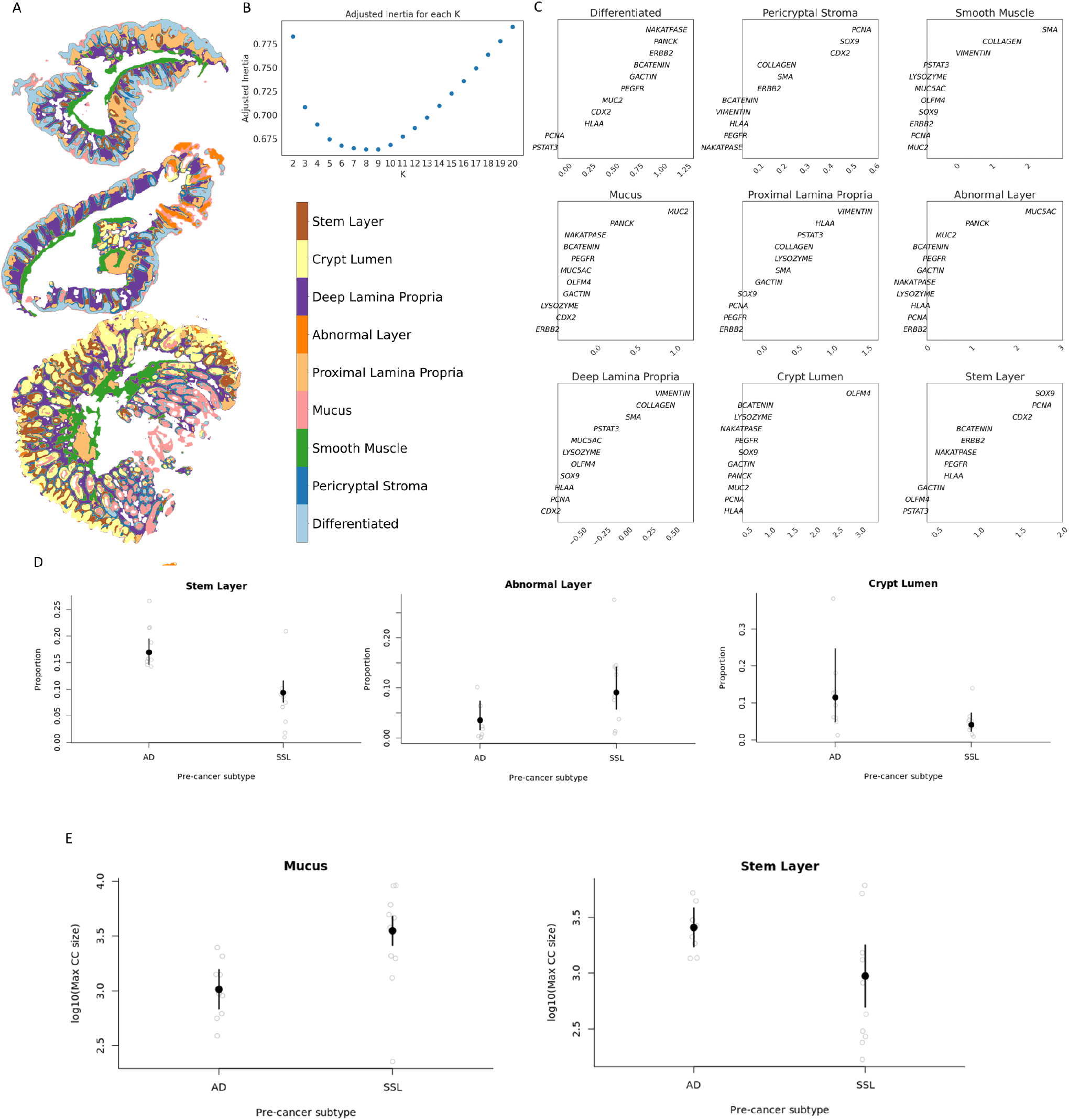
MILWRM tissue domains describe the molecular distinction between human colon adenoma pre-cancer subtypes. (A) Three representative colon mIF images with labelled tissue domains (*α* = 0.02) (B) Estimated number of tissue domains in Adjusted inertia plot (C) Domain profile describing marker composition of each tissue domain (D) GEE model results for association between tissue domains and pre-cancer subtype (E) GEE model results for association between size of connected components in tissue domains and pre-cancer subtype.

We then asked whether the two pre-cancer subtypes, AD and SER, have any differences in organization of MILWRM tissue domains. We used generalized estimating equations (GEE) to model the association of MILWRM tissue domain proportions with tumor type and found a significant association between MILWRM proportions for crypt lumen, abnormal, and stem classes (Table 1; Figure 3D). Specifically, ADs were associated with higher proportions of pixels labelled as stem and crypt lumen classes, consistent with their characteristic increased stemness driven by WNT-signaling (14, 19). In contrast, serrated polyps were associated with increased pixel proportions of the abnormal class marked by MUC5AC; MUC5AC is a foregut endoderm mucin characteristic of metaplasia associated with serrated polyps (49). AD arises from stem cell expansion which inevitably fill the entirety of abnormal crypts (19). Thus, we also hypothesized that the stem MILWRM domain will be significantly more connected compared with SER tissues. We again used GEE to estimate the population average effect of pre-cancer subtype on MILWRM the maximum size of tissue-connected components (Table 2; Figure 3E) and found a significant association between connectedness of stem and mucus tissue domains and pre-cancer type. Stem domain was expectedly more connected in AD subtype whereas higher connectedness in mucus domain was associated with SER pre-cancer type. ADs have defects in differentiation of goblet cells that inherently depletes the mucus layer (16, 22, 37, 46, 47, 62). This aligns with association of AD with decreased connected mucus components (Figure 3E). There was no such association observed for connectedness of the abnormal MUC5AC+ domain since it comprises sporadic abnormal cells associated with secretion. These results align with recent atlas results demonstrating that ADs arose from stem cell expansion and serrated polyps from pyloric metaplasia (19, 49).

### MILWRM applied to spatial transcriptomics reliably identify tissue domains across different mouse brain cross-sections

We applied MILWRM to a publicly available 10X Genomics Visium dataset comprising seven mouse brain samples including three coronal, two sagittal anterior, and two sagittal posterior slices (Figure 4A; Figure S5A-B) (1–7). We used the penalty parameter 0.02 for high resolution domain detection similar to the above for mIF data to distinguish functionally relevant brain regions (Figure 3B). We identified thirteen tissue domains in the brain ST data, and manually annotated them using histological information with a reference atlas from the Allen Brain Institute (Figure 4A – middle column) (54). Confidence score overlays demonstrate high quality and robust identification of most tissue domains (Figure 4A – right column). Notably, MILWRM identified consensus domains despite differences in the orientations and cuts of brain slices. For example, MILWRM was able to capture tissue domains that are unique only to certain slices, such as cerebellum specific to sagittal-posterior cut, as well as domains with diverse shapes and sizes due to orientation differences, such as the striatum that is small in the coronal slice, large in the sagittal-anterior slice, and absent in the sagittal-posterior cut (Figure 4A-B). The MILWRM model captures approximately 70% of variance in ST data, similar to mIF results (Figure 4C; Figure S5C).

**Fig. 4.**
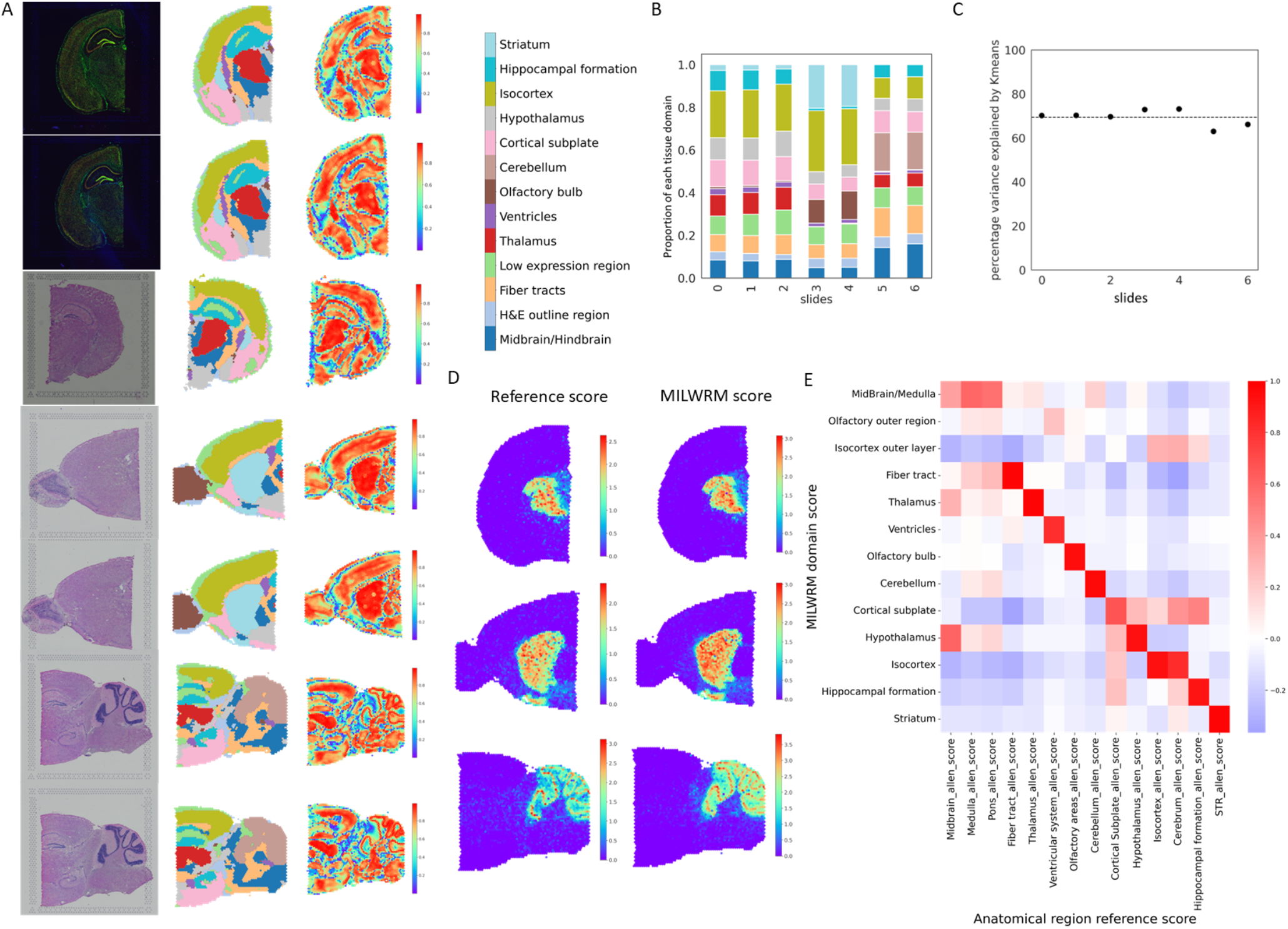
MILWRM detects consensus tissue domains in ST data from different mouse brain cross-sections. (A) MILWRM detected tissue domains (at *α* = 0.02, middle) in mouse brain ST data (H&E, left) and confidence scores (right) (B) Proportion of tissue domains in slides (C) Percentage variance explained by Kmeans (D) Reference (left) and MILWRM scores (right) for Thalamus, Striatum and Cerebellum (top to bottom respectively) (E) Overall correlation between MILWRM and reference scores for each tissue domain and anatomical region across all spots.

After histologically annotating the tissue domains using the reference atlas, we evaluated the ability of MILWRM to identify known domain distinguishing genes for potential use in unsupervised analysis. To achieve that, we curated a reference gene list for the corresponding histological regions from differential expression lists available at the Allen Brain Atlas obtained from ISH data 52. Reference lists for histological regions not available in Allen Brain Atlas were curated from the molecular atlas of mouse brain (43), which was generated from ST data. We first compared MILWRM domain-specific gene lists to curated reference gene lists for the corresponding histological regions. To validate the reference gene lists, we computed a signature score for each curated reference gene list per brain region, and then overlaid these signatures onto ST data. Reference gene signatures were expectedly highly specific to their respective brain regions (Figure 4D – left column; Figure S6A-G). In a similar vein, we also computed and overlaid MILWRM domain-specific signature scores and found that they were highly specific and accurately marked each histological brain region (Figure 4D – right column; Figure S7A-G). To quantify the performance between reference gene signatures and MILWRM signatures, we calculated a spot-by-spot correlation of the two sets of signature scores across all slides. High correlation between the MIL-WRM and reference scores was observed on a brain regionspecific basis (Figure 4E). These results illustrate that the MILWRM approach can be effectively applied to genomescale ST data for extracting tissue domain-specific molecular information.

### MILWRM performs favorably when compared to SpaGCN

While there is a paucity of methods to identify and enumerate spatial domains across samples, we compared MILWRM to recently published SpaGCN, which is one of the only algorithms that can detect spatial domains on ST data over multiple samples (30). MILWRM and SpaGCN were performed on five brain ST slides analyzed above with effectively the same resolution (MILWRM *α* = 0.01, SpaGCN res = 0.52, p = 0.5) (Figure 5A). MILWRM was able to further sub-classify previously detected tissue domains into subregions. For instance, isocortex was divided into three additional layers and cerebellum into two layers, which corresponded to brain anatomy in the reference atlas (Figure 5A - middle column, Figure 5B). These finer sub-classifications were detected across multiple slices by MILWRM. In contrast, SpaGCN was unable to detect consensus spatial domains across all slides. Only common domains detected in similarly oriented cuts were identified, whereas the same domain across uniquely sliced slides were identified separately (Figure 5A – right column). SpaGCN, when performed at varying resolutions, was also unable to identify consensus domains across replicates slides (Figure 5C - Colored arrows). Although a consensus can be reached by searching for the right parameters, it is not consistent for all domains. These results further illustrate the ability of MILWRM as one of the only algorithms to robustly identify consensus tissue domains across slides with pixel information.

**Fig. 5.**
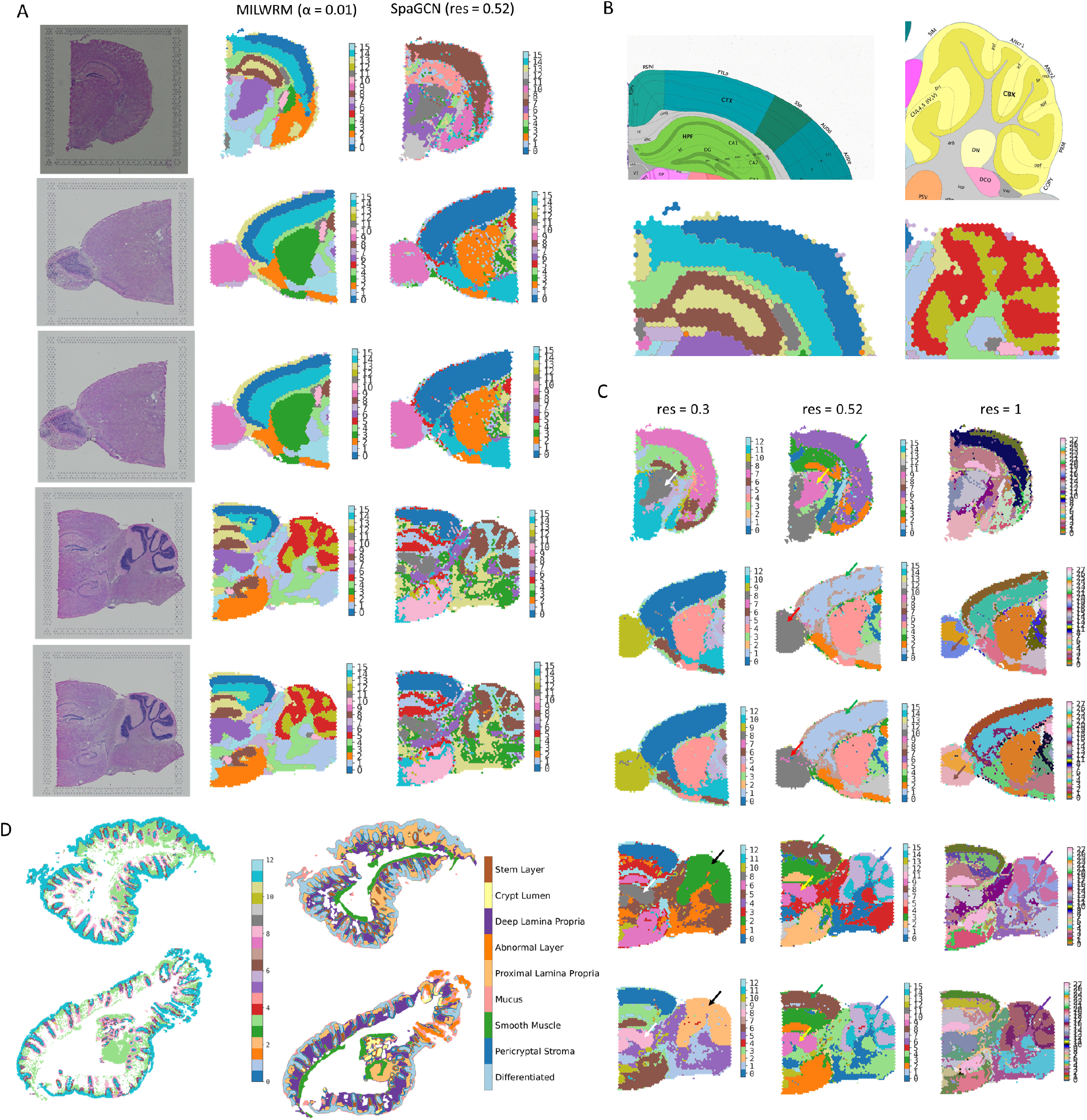
MILWRM performs better than SpaGCN. (A) MILWRM (middle) and SpaGCN (right) detected tissue domains in mouse brain ST data (B) Stratified layers in Isocortex and Cerebellum from Allen brain atlas (top) and MILWRM domains (bottom) (C) SpaGCN domains at different resolutions (res 0.3, 0.52 and 1), colored arrows point the domains that should be consensus (D) SpaGCN (left) and MILWRM domains (right) in two colon mIF specimens

While SpaGCN was only designed for ST data, we still compared its performance on mIF data, as there are currently no existing algorithms for domain detection for more than one sample in mIF data. To enable SpaGCN which only works on low resolution ST data, we performed SpaGCN spatial detection on two mIF slides downsampled to 1/32 resolution with p = 0.5, res = 0.5. While the entire tissue in pixel space was classified into MILWRM domains, there were missing tissue portions in domains detected by SpaGCN (Figure 5D – left column). Additionally, SpaGCN detected twelve consensus spatial domains across MxIF slides, but many of these domains were spurious; only three domains predominantly represented real tissue regions. Furthermore, SpaGCN was unable to capture tissue domains that classify disease tissue subtypes (SER versus normal, for example), which was apparent in the MILWRM analysis across the two slides (Figure 5D – right column). Overall, MILWRM offers a robust and flexible approach for consensus domain identification across specimens that is generally applicable to different spatial molecular data types.

## Discussion

Pixel-based tissue domain detection forms the basis of the top-down approach to spatial data analysis. Current methods of tissue domain detection are either based on a bottom-up approach, that is, building cellular neighborhoods using segmented single-cell data (20, 34, 59) and/or lack scalability across samples (8, 30, 55, 63). Here, we addressed this gap by developing MILWRM, an algorithm to detect spatial domains across samples through a top-down, pixel-based approach. We demonstrated applicability of MILWRM to find relevant biological phenotypes in multiple data modalities (MxIF and ST) in an unsupervised way without manual thresholding and annotation.

An important demonstration of MILWRM is its ability to discern organizational differences in tissue domains related to disease subtypes. While abnormal tissues can be distinguished from normal tissues within a slide using other methods, MILWRM application across slides has significant value. There are specimens that are completely composed of abnormal tissues. In those circumstances, comparison between specimens (normal vs abnormal) is the only way to distinguish between disease states. In addition, MILWRM’s ability to identify consensus tissue domains across specimens makes it possible to classify patients into disease subtypes based on tissue features. Finally, MILWRM was able to identify consensus domains and gene lists that match with organ anatomical features despite different cuts and orientations. This is important because the tissue structure from individual cuts may appear morphologically different but is functionally identical. These examples showed the real-world application of MILWRM in pathological diagnosis of disease subtypes and anatomic classification and characterization.

mIF data present additional pixel analysis obstacles. First, due to lower marker dimensionality, marker selection and management is of utmost importance. Unlike ST data where vectors of genes define programs and phenotypes, mIF phenotypes are usually defined by single markers. Highly expressed markers may mask lower expression markers if sub-optimal preprocessing is performed, thus preventing some tissue domains from being detected. Secondly, high resolution microscopy data are generally incompatible with pixel-based algorithms built for low resolution ST data, such as SpaGCN. Creation of image tiles or large-scale downsampling is needed to satisfy speed and memory requirements. In contrast to most state-of-the-art methods for pixel-based analysis that are data type-specific, MILWRM is adaptable to multiple imaging data types and is scalable to many samples. While SpaGCN is a scalable method for spatial domain detection, it failed to discern disease-specific differences in tissues from the tissue domains it identified. Additionally, the domains identified by SpaGCN in ST data were not robust since they failed to reach consensus at different clustering resolutions. Additionally, MILWRM also provides various QC metrics which can be used to assess the quality of domain detection and ability to perform tissue clustering at different levels of smoothing, downsampling, and cluster resolution.

## Methods

### Resource availability

#### Lead Contact

Request for further information and requests for resources should be directed to and will be fulfilled by the corresponding authors, Ken Lau and Simon Vandekar

#### Materials Availability

This study did not generate any new unique reagents.

#### Data and Code Availability

The package is available to be installed directly from pyPI website (https://pypi.org/project/MILWRM/). The source code for the package and code for the figures can be found on github (https://github.com/Ken-Lau-Lab/MILWRM).

### MILWRM workflow details

#### Data preprocessing

Spatial-omics data differ in their acquisition and technological artifacts across modalities, so these preprocessing steps are data type specific. It is important for the user to understand and apply methods that are reasonable for their modality before using MILWRM, otherwise results can be corrupted by batch effects.

#### A. Multiplex Immunofluorescence (mIF)

Prior to preprocessing, mIF data were scaled from uint8 (0 to 255) to float (0 to 1) and downsampled by a factor 1/16th resolution to speed computation and normalization process. There is no sacrifice in the quality of the neighborhood identification by downsampling as mIF data have subcellular spatial resolution and MILWRM is designed to identify broad tissue domains. After downsampling we created tissue masks for each image as described in mIF tissue mask generation with MILWRM. Finally, we applied image normalization at slide-level using the formula 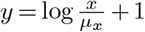, where x is the unnormalized data and *μ_x_* is the mean of non-zero pixels in the image, per marker. This normalization was a modification of an existing method evaluated in segmented mIF data. Here, we implemented the mean of non-zero pixels to accommodate channels with sparse signal intensities (27). The downsampling performed on images prior to this normalization step also aligns with the unbiased grid-based normalization framework described by Graf et al., 2022 (24). In order to incorporate broader spatial information within each pixel, after normalization, we applied gaussian smoothing. The radius of blurring can be controlled by adjusting *σ* parameter in MILWRM for mIF modality. Here, we use *σ* = 2 for smoothing.

#### B. Spatial Transcriptomics (ST)

The above-described steps differ slightly between mIF and ST modality. For ST data the first step is to reduce the dimensionality of the transcriptomics data. For the analysis shown in this paper, we used Principal Component Analysis (PCA) for dimensionally reduction, but other methods can also be used with MILWRM such as Non-Negative Matrix Factorization (NMF) (36). We used Harmony (35) to correct technical variation between the samples. As in mIF, blurring is applied to the ST slides to preserve spatial information. To perform blurring, each central spot is assigned the average value for the selected reduced components (PCs in this case) across the spots within the neighborhood of the central spot. The spatial neighborhoods are computed using the `squidpy` Python package (44). The neighborhood distance can be controlled by adjusting the `n_rings` parameter. Here, we use `n_rings` = 1 for smoothing in ST data.

#### Identification of tissue domains

The tissue domains in the data are identified across slides by performing unsupervised K-means clustering on the preprocessed data. MILWRM reduced computation time by randomly subsampling pixels for mIF modality. The fraction of pixels (default 0.2, used here) and all the spots are serialized to build the K-means model. If the dimension reduction is performed, then the input data are the PCs, otherwise the input is the batch-adjusted marker channels. Prior to performing K-means, the data are Z-normalized to ensure that the mean and variances are similar across the different channels/PCs of the input data. The k-selection for K-means is done by estimating adjusted inertia metric. Adjusted inertia is inertia weighted by a penalty parameter that controls the number of clusters (21). For MILWRM, the parameter can be adjusted to control the resolution of tissue domains identified.

After performing K-means classification, tissue domains are identified in the full dataset by assigning the tissue domain for the closest cluster centroid from the K-means model. The mean and variance computed for subsampled data is used to Z-normalize the original image data. By performing K-means model estimation in the subsample, MILWRM can reduce computational demand for mIF modality. Kmeans is performed on entire dataset in ST modality.

#### Quality control and tissue labeling

Once the regions are identified it is useful to label the tissue domains based on their marker expression profile and assess the quality of clustering. The cluster centroids for each tissue domain are plotted in marker or PCA component space to label the tissue cluster based on its expression profile. The centroids can also be plotted in gene space for ST modality or other dimensionally reduced components. The quality of clustering is assessed at the whole slide and pixel levels. To assess the whole slide fit, we compute the variance explained and mean square error within each slide. These metrics allow the user to flag slides for manual review where the overall fit might be bad. We also compute pixel-level confidence score using the formula 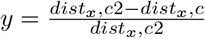 where `dist` is the Euclidean distance between pixel or spot, `x`, assigned centroid, `c`, and the second closest centroid `c`. The confidence scores take values between zero and one where higher values indicate smaller distance between the assigned centroid and closest centroid thus, better fit. This metric is a fast simplification of the Silhouette index.

#### mIF tissue mask generation with MILWRM

MILWRM has a designated function to perform creation of tissue masks through the MILWRM pipeline described above. Each preprocessing step is performed on individual images including log normalization and smoothing with a gaussian filter (*σ* = 2). Finally, the mask is created using Kmeans clustering with n = 2. The Kmeans cluster centers are then z-normalized and the cluster center with a mean smaller or equal to zero is set as background.

#### Imaging data, acquisition, and basic image processing

The mIF data were generated for the Human Tumor Atlas Network (HTAN) consisting of human normal colon and different colonic pre-cancer subtypes (conventional adenomas - AD and serrated polyps - SER) (19). These data comprised multichannel fluorescent images from 37 biospecimen consisting of tissues with different morphologies and pathological classification, as confirmed by 2 pathologists (Supplementary Table 1). Cyclical antibody staining, detection, and dye inactivation was performed as described previously (23). In brief, fluorescent images were acquired at 200x magnification on a GE In Cell Analyzer 2500 using the Cell DIVE platform. Exposure times were determined for each antibody. Dye inactivation was accomplished with an alkaline peroxide solution, and background images were collected after each round of staining to ensure fluorophore inactivation. Staining sequence, conditions, and exposure times are as described in (19). Following acquisition, images were processed as described (41). Briefly, DAPI images for each round were registered to a common baseline, and autofluorescence in staining rounds was removed by subtracting the previous background image for each position.

#### Method evaluation and statistical analysis

In order to assess the sensitivity of MILWRM regions to biological differences between precancer subtypes we computed tissue proportions and connected component statistics for each tissue domain within the tumor region of each image and used generalized estimating equations (GEEs) to model how these variables were associated with precancer subtypes. Connected components were estimated for each image in Python using the label function in `scipy.ndimage.measurements` module. For the tissue proportions, we modeled each tissue proportion separately using a binomial family model assuming that images from the same slide had an exchangeable correlation structure. We modeled the maximum connected component size in order to quantify how the size and connectedness of different tissue domains differed across precancer subtypes. In these analyses, we used log transformation in a gaussian family model with a log transformation on the maximum connected component size and included log of the total tissue volume as a covariate. In all models, we weighted each region by its total image size so that results were not affected by noisy estimates from smaller images. Statistical analyses were performed in R using the `geepack` package (26). We performed plot all results with unadjusted significant p-values and report adjusted p-values using the Benjamini-Hochberg procedure and a robust effect size index (15, 57) (Tables 1 and 2).

#### Tissue domain signature scores for ST data

The manual annotation for tissue domains in ST data were verified by generating signature gene scores specific to each brain region. For this purpose, we extracted differentially expressed genes from Allen brain atlas for all available brain regions and molecular atlas of adult mouse brain for fiber tract and ventricles. MILWRM also identified a set of genes for each tissue domain. We computed a score for both reference signature set and MILWRM gene set using `scanpy` (60).

