## Supplementary Figures for "Consensus tissue domain detection in spatial multi-omics data using MILWRM"

**Supplementary Table 1. Summary table for human colonic adenoma sample metadata**

| Batch name | Slide region | Tissue category | Broad precancer type |
|---|---|---|---|
| HTA11_10167_0000_01_01 | HTA11_10167_0000_01_01_region_001 | normal | SSL |
| HTA11_10167_0000_01_01 | HTA11_10167_0000_01_01_region_002 | normal | SSL |
| HTA11_10167_0000_01_01 | HTA11_10167_0000_01_01_region_003 | Tumor | SSL |
| HTA11_10623_0000_01_01 | HTA11_10623_0000_01_01_region_001 | normal | AD |
| HTA11_10623_0000_01_01 | HTA11_10623_0000_01_01_region_002 | normal | AD |
| HTA11_10623_0000_01_01 | HTA11_10623_0000_01_01_region_003 | normal | AD |
| HTA11_10623_0000_01_01 | HTA11_10623_0000_01_01_region_004 | Tumor | AD |
| HTA11_10623_0000_01_01 | HTA11_10623_0000_01_01_region_005 | normal | AD |
| HTA11_10623_0000_01_01 | HTA11_10623_0000_01_01_region_006 | normal | AD |
| HTA11_10711_0000_01_01 | HTA11_10711_0000_01_01_region_001 | Tumor | AD |
| HTA11_4255_0000_02_02 | HTA11_4255_0000_02_02_region_001 | Tumor | SSL |
| HTA11_4255_0000_02_02 | HTA11_4255_0000_02_02_region_002 | normal | SSL |
| HTA11_6298_0000_04A_03 | HTA11_6298_0000_04A_03_region_001 | Tumor | AD |
| HTA11_6298_0000_04A_03 | HTA11_6298_0000_04A_03_region_002 | Tumor | AD |
| HTA11_6298_0000_04A_03 | HTA11_6298_0000_04A_03_region_003 | normal | AD |
| HTA11_6801_0000_01_01 | HTA11_6801_0000_01_01_region_001 | Tumor | SSL |
| HTA11_7179_0000_02_02 | HTA11_7179_0000_02_02_region_001 | Tumor | AD |
| HTA11_7862_0000_02_02 | HTA11_7862_0000_02_02_region_001 | Tumor | AD |
| HTA11_7862_0000_02_02 | HTA11_7862_0000_02_02_region_002 | normal | AD |
| HTA11_7956_0000_02_05 | HTA11_7956_0000_02_05_region_001 | normal | SSL |
| HTA11_7956_0000_02_05 | HTA11_7956_0000_02_05_region_002 | Tumor | SSL |
| HTA11_7956_0000_02_05 | HTA11_7956_0000_02_05_region_003 | Tumor | SSL |
| HTA11_8099_0000_02_01 | HTA11_8099_0000_02_01_region_001 | normal | SSL |
| HTA11_8099_0000_02_01 | HTA11_8099_0000_02_01_region_002 | normal | SSL |
| HTA11_8099_0000_02_01 | HTA11_8099_0000_02_01_region_003 | normal | SSL |
| HTA11_8099_0000_02_01 | HTA11_8099_0000_02_01_region_004 | normal | SSL |
| HTA11_8099_0000_02_01 | HTA11_8099_0000_02_01_region_005 | Tumor | SSL |
| HTA11_8099_0000_02_01 | HTA11_8099_0000_02_01_region_006 | normal | SSL |
| HTA11_8622_0000_01E_01 | HTA11_8622_0000_01E_01_region_001 | normal | SSL |
| HTA11_8622_0000_01E_01 | HTA11_8622_0000_01E_01_region_002 | Tumor | SSL |
| HTA11_8622_0000_01E_01 | HTA11_8622_0000_01E_01_region_003 | Tumor | SSL |
| HTA11_8622_0000_01E_01 | HTA11_8622_0000_01E_01_region_004 | Tumor | SSL |
| HTA11_8622_0000_01E_01 | HTA11_8622_0000_01E_01_region_005 | normal | SSL |
| HTA11_866_0000_02_03 | HTA11_866_0000_02_03_region_001 | Tumor | AD |
| HTA11_8920_0000_02_02 | HTA11_8920_0000_02_02_region_001 | Tumor | SSL |
| HTA11_9341_0000_01A_01 | HTA11_9341_0000_01A_01_region_001 | Tumor | SSL |
| HTA11_9408_0000_02A_05 | HTA11_9408_0000_02A_05_region_001 | Tumor | AD |
| HTA11_9408_0000_02A_05 | HTA11_9408_0000_02A_05_region_002 | Tumor | AD |

**Table S2. Coefficient summary table for association between size of tissue domain and pre-cancer subtype**

| Pixel size of tissue domain | Estimate | SE | Chi-square | p-value | RESI | $p_{FDR}$ |
|---|---|---|---|---|---|---|
| Differentiated tissue domain | 0.412 | 2.12e-01 | 3.774 | 0.052 | 0.430 | 0.130 |
| Pericryptal Stroma | 0.240 | 1.31e-01 | 3.366 | 0.067 | 0.397 | 0.133 |
| Smooth Muscle | -0.341 | 2.83e-01 | 1.449 | 0.229 | 0.173 | 0.381 |
| Mucus | 0.036 | 1.68e-01 | 0.046 | 0.830 | 0.000 | 0.922 |
| Deep Lamina Propria | 0.010 | 2.10e-01 | 0.002 | 0.962 | 0.000 | 0.962 |
| Abnormal Layer | 0.996 | 4.68e-01 | 4.534 | 0.033 | 0.485 | 0.130 |
| Proximal Lamina Propria | 0.145 | 1.87e-01 | 0.604 | 0.437 | 0.000 | 0.581 |
| Crypt Lumen | -1.116 | 5.65e-01 | 3.904 | 0.048 | 0.440 | 0.130 |
| Stem Layer | -0.680 | 1.48e-01 | 21.220 | 0.000 | 1.161 | 0.000 |
| Total | 50593.575 | 6.92e+04 | 0.535 | 0.464 | 0.000 | 0.581 |

**Table S3. Coefficient summary table for association between size of maximum connected component of each tissue domain and pre-cancer type**

| Maximum pixel size of connected components in tissue domain | Estimate | SE | Chi-square | p-value | RESI | $p_{FDR}$ |
|---|---|---|---|---|---|---|
| Differentiated tissue domain | 0.045 | 0.178 | 0.065 | 0.798 | 0.000 | 0.898 |
| Pericryptal Stroma | 0.012 | 0.136 | 0.008 | 0.929 | 0.000 | 0.929 |
| Smooth Muscle | -0.164 | 0.144 | 1.299 | 0.254 | 0.141 | 0.357 |
| Mucus | 0.535 | 0.105 | 25.849 | 0.000 | 1.287 | 0.000 |
| Deep Lamina Propria | 0.114 | 0.084 | 1.836 | 0.175 | 0.236 | 0.357 |
| Abnormal Layer | 0.275 | 0.254 | 1.177 | 0.278 | 0.109 | 0.357 |
| Proximal Lamina Propria | -0.142 | 0.112 | 1.595 | 0.207 | 0.199 | 0.357 |
| Crypt Lumen | -0.262 | 0.193 | 1.850 | 0.174 | 0.238 | 0.357 |
| Stem Layer | -0.435 | 0.175 | 6.164 | 0.013 | 0.587 | 0.059 |

A

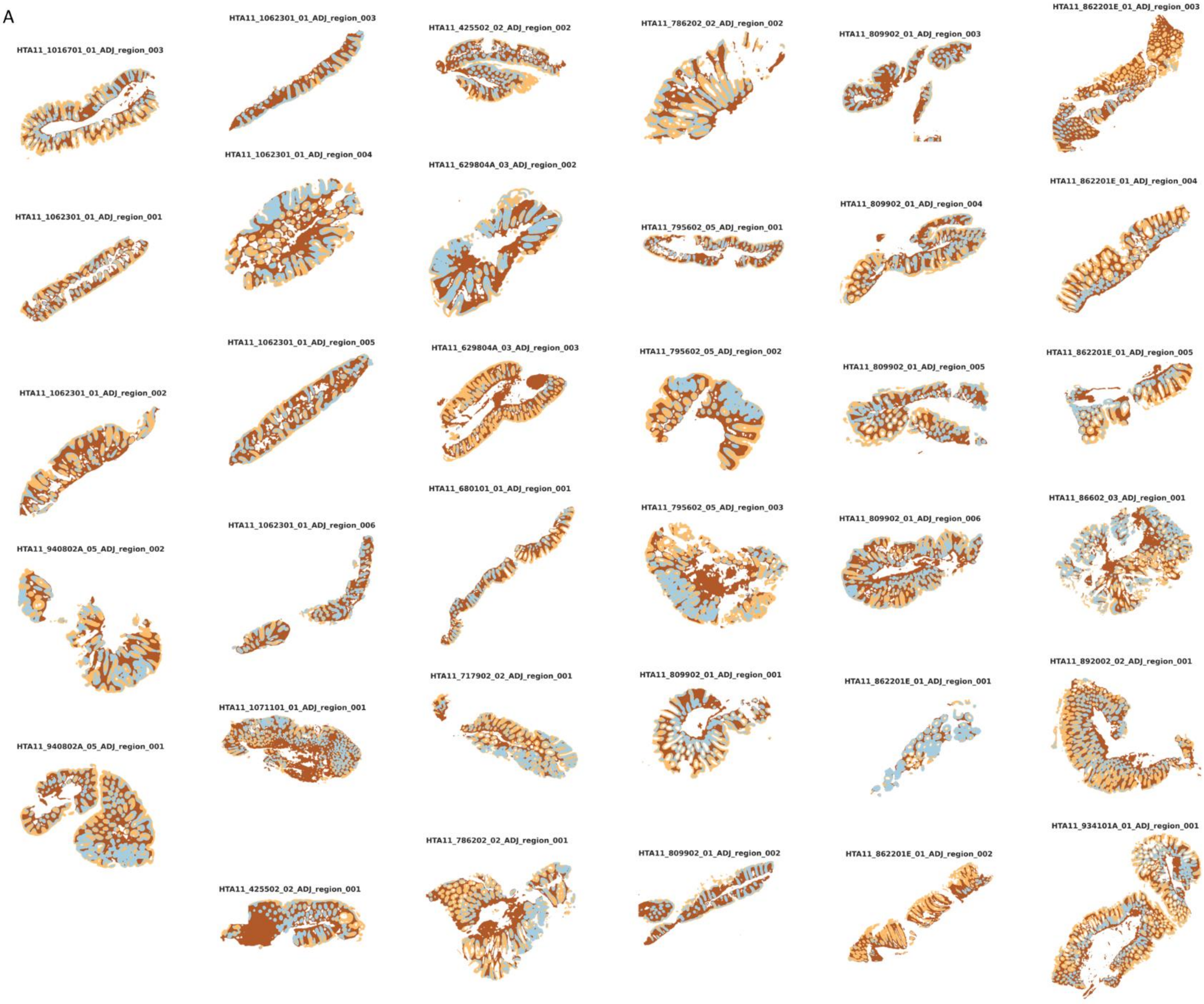

HTA11_1016701_01_ADJ_region_003
HTA11_1062301_01_ADJ_region_003
HTA11_425502_02_ADJ_region_002
HTA11_786202_02_ADJ_region_002
HTA11_809902_01_ADJ_region_003
HTA11_862201E_01_ADJ_region_003
HTA11_1062301_01_ADJ_region_004
HTA11_629804A_03_ADJ_region_002
HTA11_862201E_01_ADJ_region_004
HTA11_809902_01_ADJ_region_004
HTA11_1062301_01_ADJ_region_001
HTA11_795602_05_ADJ_region_001
HTA11_1062301_01_ADJ_region_005
HTA11_629804A_03_ADJ_region_003
HTA11_795602_05_ADJ_region_002
HTA11_862201E_01_ADJ_region_005
HTA11_809902_01_ADJ_region_005
HTA11_1062301_01_ADJ_region_002
HTA11_680101_01_ADJ_region_001
HTA11_795602_05_ADJ_region_003
HTA11_86602_03_ADJ_region_001
HTA11_809902_01_ADJ_region_006
HTA11_1062301_01_ADJ_region_006
HTA11_940802A_05_ADJ_region_002
HTA11_717902_02_ADJ_region_001
HTA11_809902_01_ADJ_region_001
HTA11_892002_02_ADJ_region_001
HTA11_862201E_01_ADJ_region_001
HTA11_1071101_01_ADJ_region_001
HTA11_940802A_05_ADJ_region_001
HTA11_786202_02_ADJ_region_001
HTA11_934101A_01_ADJ_region_001
HTA11_809902_01_ADJ_region_002
HTA11_862201E_01_ADJ_region_002
HTA11_425502_02_ADJ_region_001


**Figure S1**

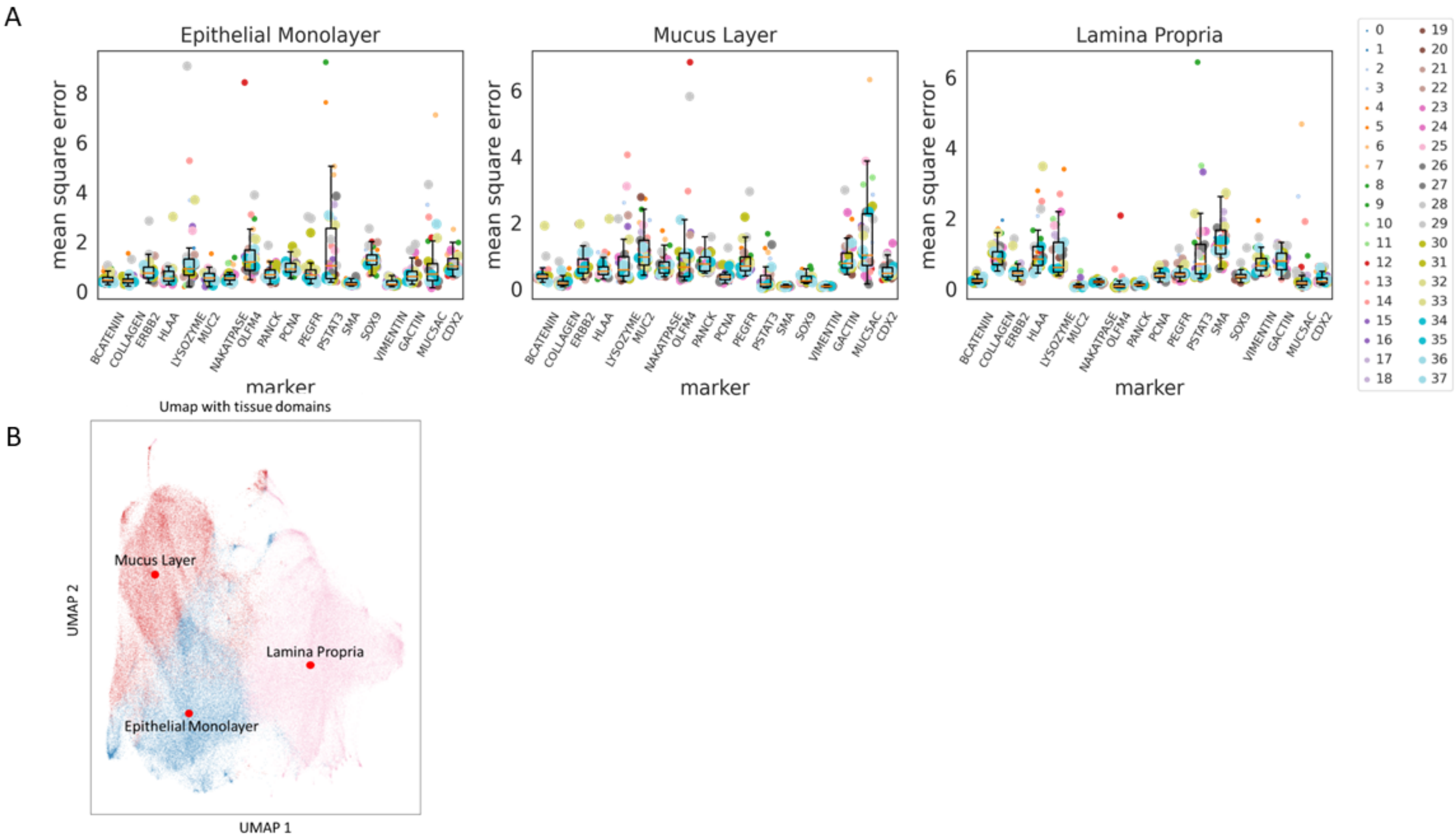
A
Epithelial Monolayer
Mucus Layer
Lamina Propria
mean square error
marker
BCATENIN
COLLAGEN
ERBB2
HLAA
LYSOZYME
MUC2
NAKATPASE
OLFM4
PANCK
PCNA
PEGFR
PSTAT3
SMA
SOX9
VIMENTIN
GACTIN
MUC5AC
CDX2
B
Umap with tissue domains
Mucus Layer
Lamina Propria
Epithelial Monolayer
UMAP 1
UMAP 2

**Figure S2**

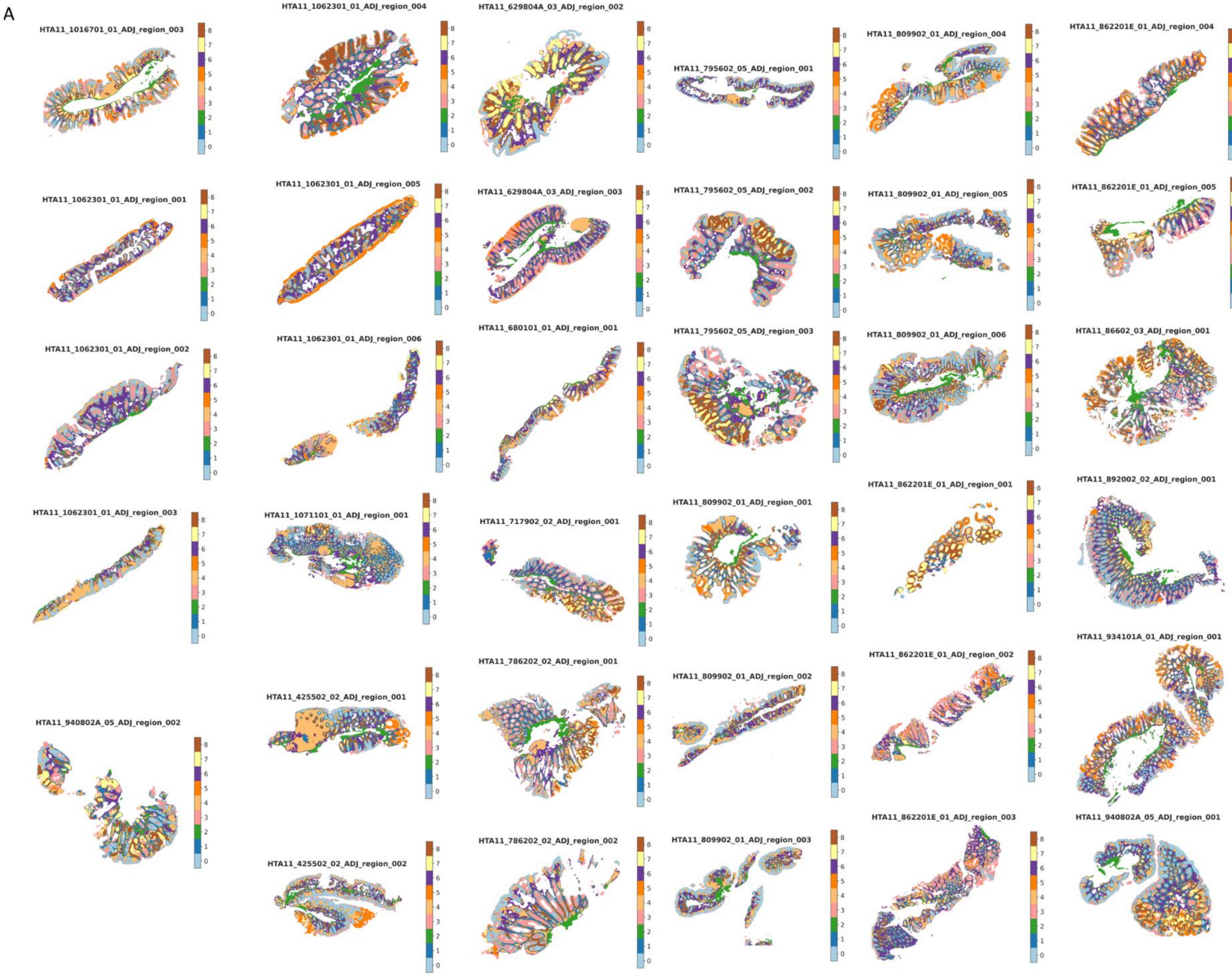
A
HTA11_1016701_01_ADJ_region_003
HTA11_1062301_01_ADJ_region_004
HTA11_629804A_03_ADJ_region_002
HTA11_795602_05_ADJ_region_001
HTA11_809902_01_ADJ_region_004
HTA11_862201E_01_ADJ_region_004
HTA11_1062301_01_ADJ_region_001
HTA11_1062301_01_ADJ_region_005
HTA11_629804A_03_ADJ_region_003
HTA11_795602_05_ADJ_region_002
HTA11_809902_01_ADJ_region_005
HTA11_862201E_01_ADJ_region_005
HTA11_1062301_01_ADJ_region_002
HTA11_1062301_01_ADJ_region_006
HTA11_680101_01_ADJ_region_001
HTA11_795602_05_ADJ_region_003
HTA11_809902_01_ADJ_region_006
HTA11_86602_03_ADJ_region_001
HTA11_1062301_01_ADJ_region_003
HTA11_1071101_01_ADJ_region_001
HTA11_717902_02_ADJ_region_001
HTA11_809902_01_ADJ_region_001
HTA11_862201E_01_ADJ_region_001
HTA11_892002_02_ADJ_region_001
HTA11_940802A_05_ADJ_region_002
HTA11_425502_02_ADJ_region_001
HTA11_786202_02_ADJ_region_001
HTA11_809902_01_ADJ_region_002
HTA11_862201E_01_ADJ_region_002
HTA11_934101A_01_ADJ_region_001
HTA11_425502_02_ADJ_region_002
HTA11_786202_02_ADJ_region_002
HTA11_809902_01_ADJ_region_003
HTA11_862201E_01_ADJ_region_003
HTA11_940802A_05_ADJ_region_001

**Figure S3**

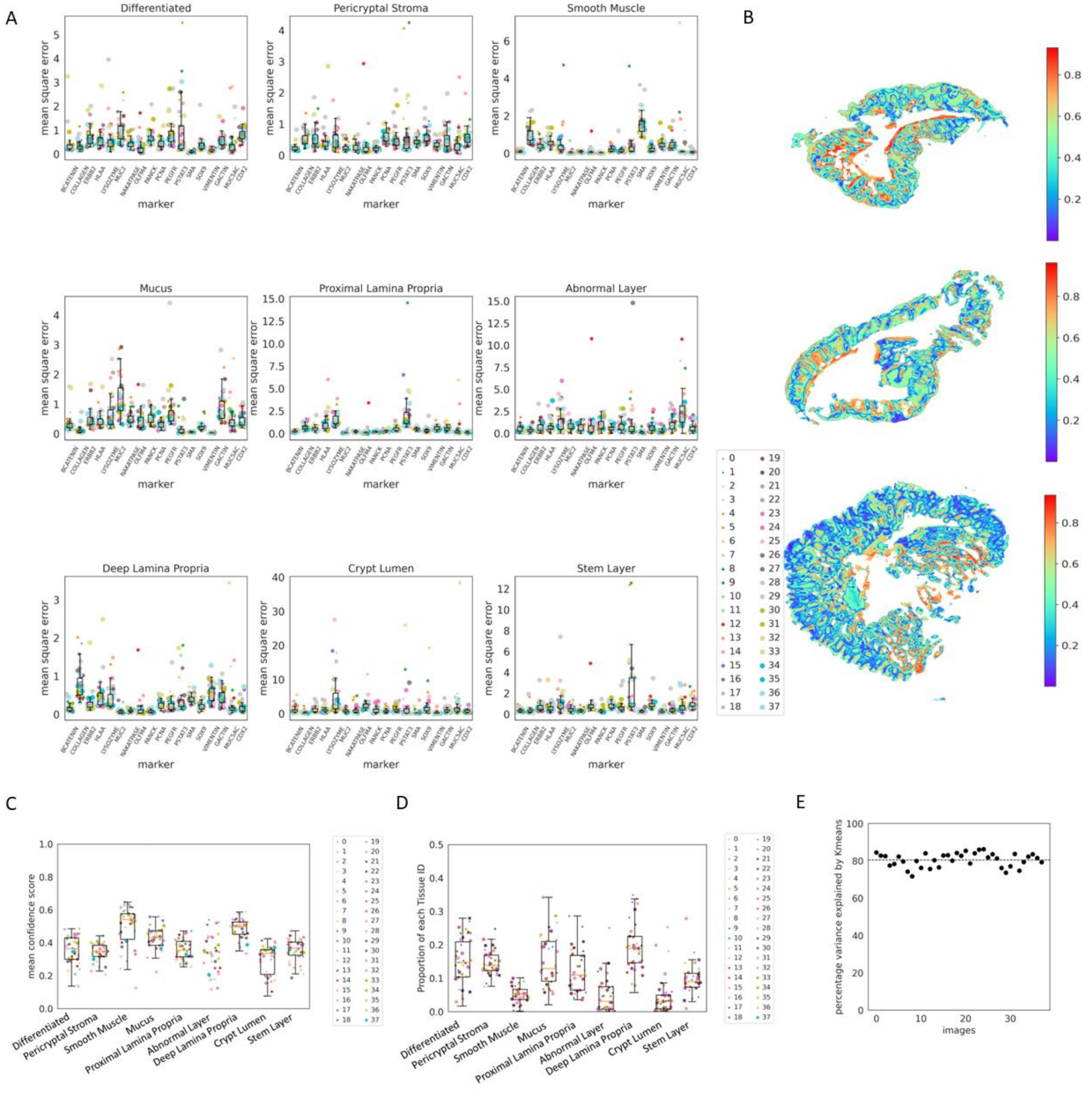
A
Differentiated
Pericryptal Stroma
Smooth Muscle
Mucus
Proximal Lamina Propria
Abnormal Layer
Deep Lamina Propria
Crypt Lumen
Stem Layer
mean square error
marker
BCATENIN
COLLAGEN
ERBB2
HLAA
LYSOZYME
MUC2
NAKATPASE
OLFM4
PANCK
PCNA
PEGFR
PSTAT3
SMA
SOX9
VIMENTIN
GACTIN
MUC5AC
CDX2
B
C
mean confidence score
D
Proportion of each Tissue ID
E
percentage variance explained by Kmeans
images

**Figure S4**

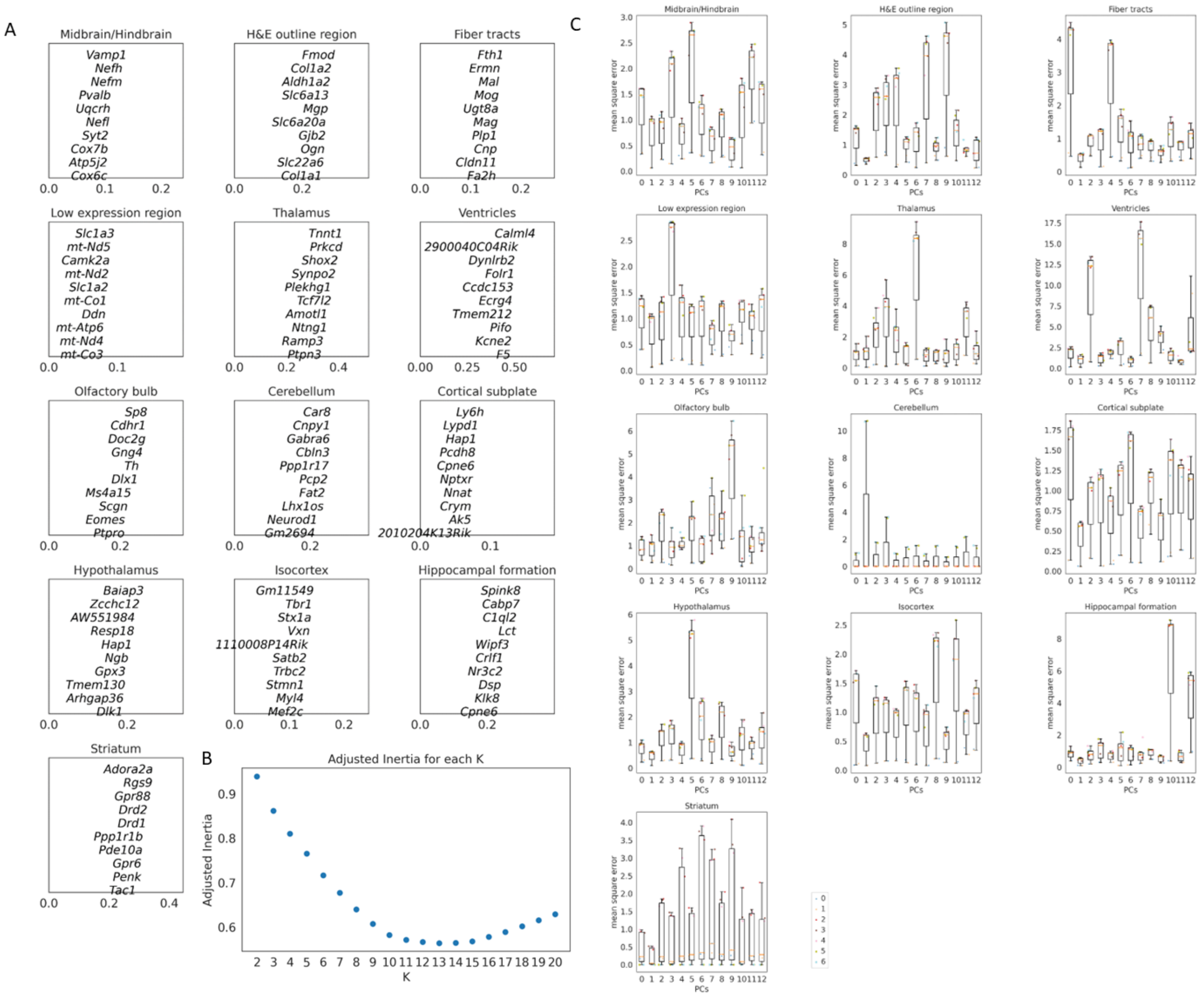
A
Midbrain/Hindbrain
Vamp1
Nefh
Nefm
Pvalb
Uqcrh
Nefl
Syt2
Cox7b
Atp5j2
Cox6c
H&E outline region
Fmod
Col1a2
Aldh1a2
Slc6a13
Mgp
Slc6a20a
Gjb2
Ogn
Slc22a6
Col1a1
Fiber tracts
Fth1
Ermn
Mal
Mog
Ugt8a
Mag
Plp1
Cnp
Cldn11
Fa2h
Low expression region
Slc1a3
mt-Nd5
Camk2a
mt-Nd2
Slc1a2
mt-Co1
Ddn
mt-Atp6
mt-Nd4
mt-Co3
Thalamus
Tnnt1
Prkcd
Shox2
Synpo2
Plekhg1
Tcf7l2
Amotl1
Ntng1
Ramp3
Ptpn3
Ventricles
Calml4
2900040C04Rik
Dynlrb2
Folr1
Ccdc153
Ecrg4
Tmem212
Pifo
Kcne2
F5
Olfactory bulb
Sp8
Cdhr1
Doc2g
Gng4
Th
Dlx1
Ms4a15
Scgn
Eomes
Ptpro
Cerebellum
Car8
Cnpy1
Gabra6
Cbln3
Ppp1r17
Pcp2
Fat2
Lhx1os
Neurod1
Gm2694
Cortical subplate
Ly6h
Lypd1
Hap1
Pcdh8
Cpne6
Nptxr
Nnat
Crym
Ak5
2010204K13Rik
Hypothalamus
Baiap3
Zcchc12
AW551984
Resp18
Hap1
Ngb
Gpx3
Tmem130
Arhgap36
Dlk1
Isocortex
Gm11549
Tbr1
Stx1a
Vxn
1110008P14Rik
Satb2
Trbc2
Stmn1
Myl4
Mef2c
Hippocampal formation
Spink8
Cabp7
C1ql2
Lct
Wipf3
Crlf1
Nr3c2
Dsp
Klk8
Cpne6
Striatum
Adora2a
Rgs9
Gpr88
Drd2
Drd1
Ppp1r1b
Pde10a
Gpr6
Penk
Tac1
B
Adjusted Inertia for each K
Adjusted Inertia
K
C
mean square error
PCs


**Figure S5**

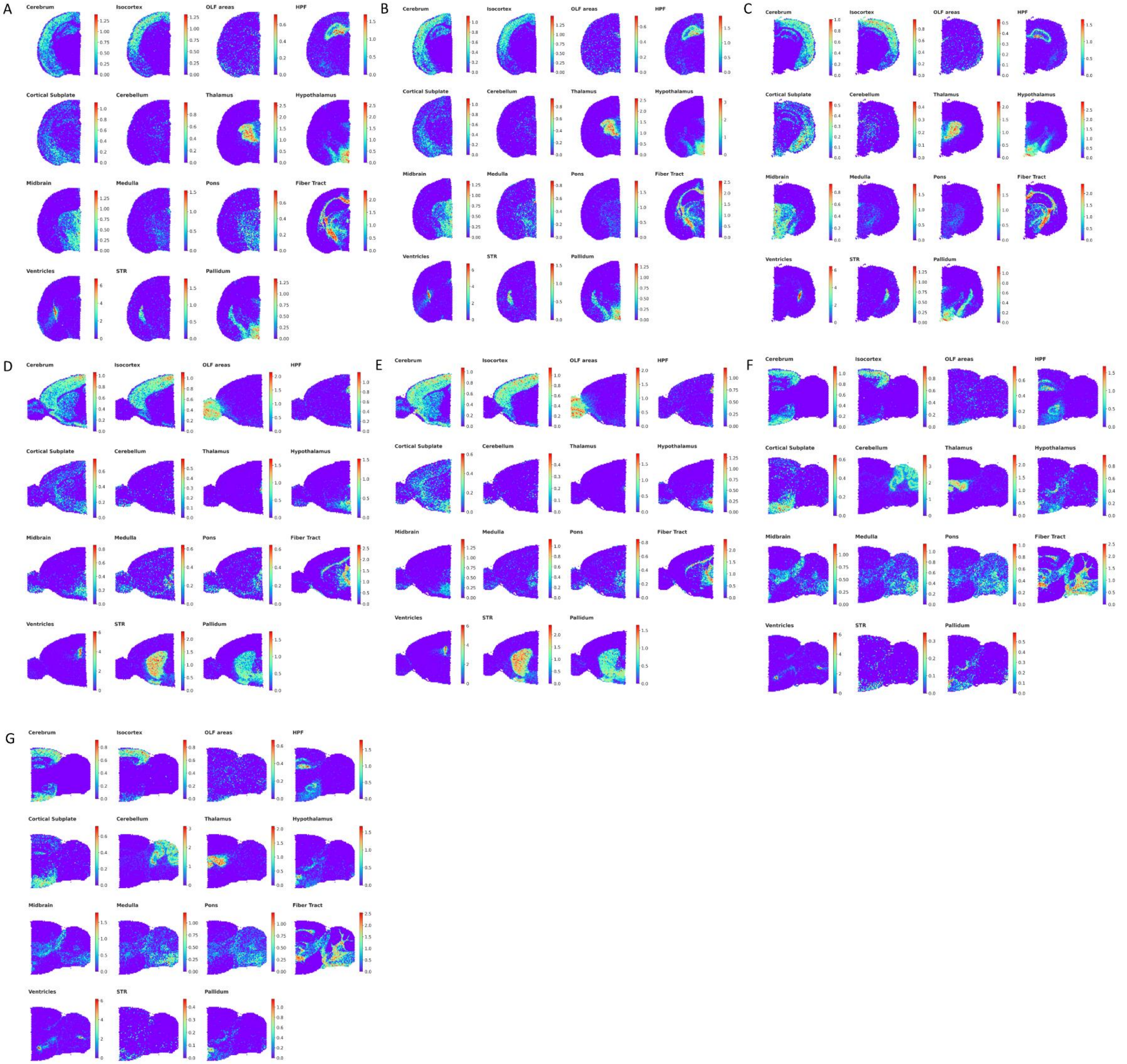

A
Cerebrum
Isocortex
OLF areas
HPF
Cortical Subplate
Cerebellum
Thalamus
Hypothalamus
Midbrain
Medulla
Pons
Fiber Tract
Ventricles
STR
Pallidum
B
Cerebrum
Isocortex
OLF areas
HPF
Cortical Subplate
Cerebellum
Thalamus
Hypothalamus
Midbrain
Medulla
Pons
Fiber Tract
Ventricles
STR
Pallidum
C
Cerebrum
Isocortex
OLF areas
HPF
Cortical Subplate
Cerebellum
Thalamus
Hypothalamus
Midbrain
Medulla
Pons
Fiber Tract
Ventricles
STR
Pallidum
D
Cerebrum
Isocortex
OLF areas
HPF
Cortical Subplate
Cerebellum
Thalamus
Hypothalamus
Midbrain
Medulla
Pons
Fiber Tract
Ventricles
STR
Pallidum
E
Cerebrum
Isocortex
OLF areas
HPF
Cortical Subplate
Cerebellum
Thalamus
Hypothalamus
Midbrain
Medulla
Pons
Fiber Tract
Ventricles
STR
Pallidum
F
Cerebrum
Isocortex
OLF areas
HPF
Cortical Subplate
Cerebellum
Thalamus
Hypothalamus
Midbrain
Medulla
Pons
Fiber Tract
Ventricles
STR
Pallidum
G
Cerebrum
Isocortex
OLF areas
HPF
Cortical Subplate
Cerebellum
Thalamus
Hypothalamus
Midbrain
Medulla
Pons
Fiber Tract
Ventricles
STR
Pallidum


**Figure S6**

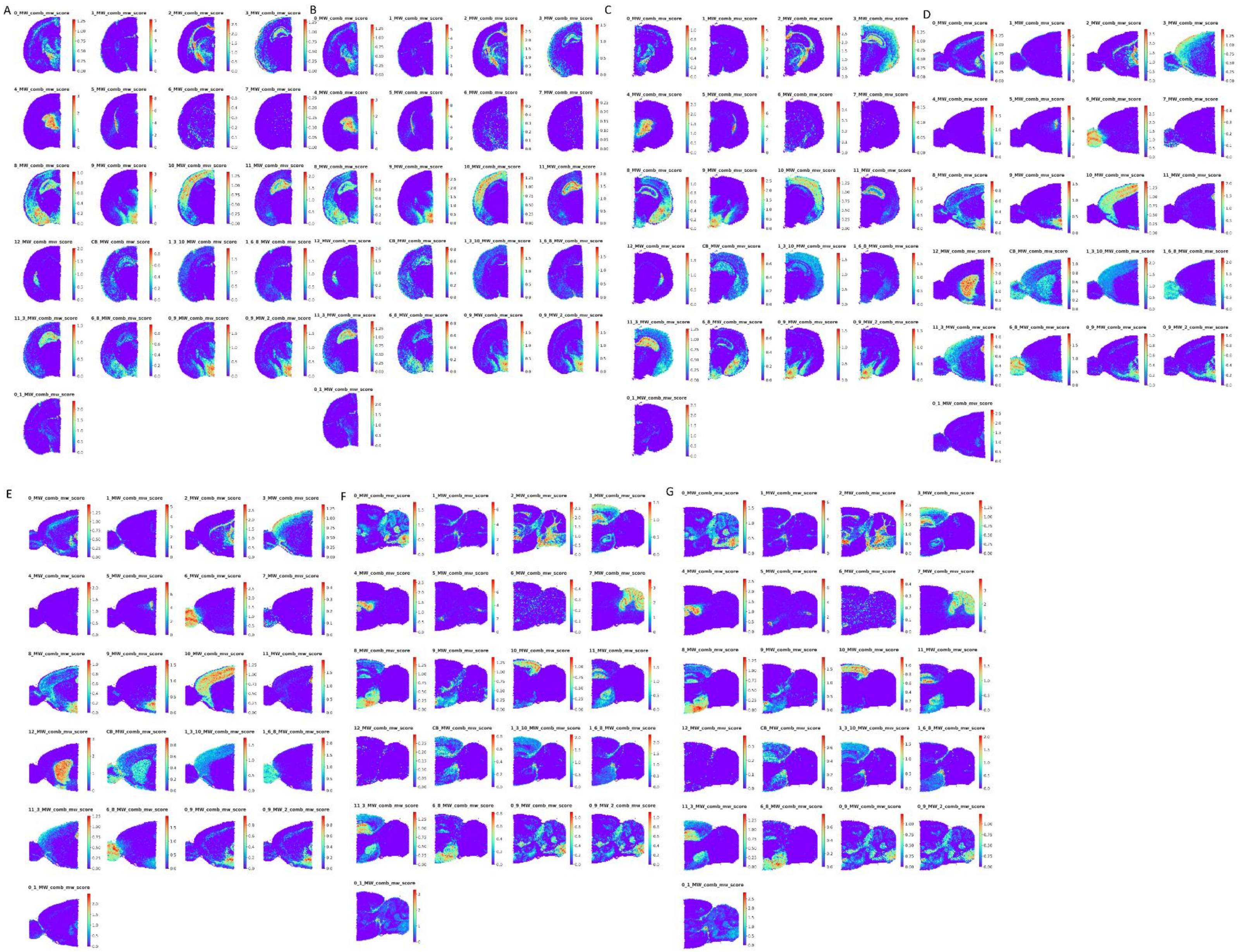

**Figure S7**

### SUPPLEMENTARY FIGURE CAPTIONS

**Figure S1. Tissue domain labels for human colonic-adenoma (alpha = 0.05).** Tissue domains for all the colonic adenoma samples (alpha = 0.05).

**Figure S2. MILWRM qc metrics for colonic MILWRM domains derived at alpha=0.05.** (A) Boxplot for mean square error within each tissue domain for each marker. Mean square error is average of Euclidean distance between each pixel intensity of that marker and Kmeans centroid for that tissue domain. Each dot with a different colored/sized combination represents a sample. Boxes represent data quartiles, and whiskers represent the interquartile ranges. (B) Umap of pixel data used for model building with tissue domain labels (color overlays) and centroids (labeled red bold dot).

**Figure S3. Tissue domain labels for human colonic-adenoma (alpha = 0.02).** Tissue domains for all the colonic adenoma samples (alpha = 0.02)

**Figure S4. MILWRM qc metrics for colonic MILWRM domains derived at alpha=0.02.** (A) Boxplot for mean square error within each tissue domain for each marker. Mean square error is average of Euclidean distance between each pixel intensity of that marker and Kmeans centroid for that tissue domain. Each dot with a different colored/sized combination represents a sample. Boxes represent data quartiles, and whiskers represent the interquartile ranges. (B) Confidence scores overlayed on three representative tissue samples. (C) Boxplot for average confidence score across all pixels in each image for each tissue domain with each dot with a different colored/sized combination representing a sample. Boxes represent data quartiles, and whiskers represent the interquartile ranges. (D) Boxplot for proportion of each tissue domain across 38 samples. (E) Scatter plot for percentage variance explained by the Kmeans model over 38 samples with each dot corresponding to a sample. Dotted line represents mean variance explained across all samples**.**

**Figure S5. Domain profile for mouse brain tissue domains and MILWRM qc metrics.** (A) Domain profile for tissue domains in mouse brain ST. (B) Estimated number of tissue domains in Adjusted Inertia plot. (C) Boxplot for mean square error within each tissue domain for each PC. Mean square error is average of Euclidean distance between PCs at each spot and Kmeans centroid for that tissue domain. Boxes represent data quartiles, and whiskers represent the interquartile ranges. Colored dots represent a sample.

**Figure S6. Reference score profiles for all mouse brain samples.** (A) Reference gene Scores from Allen Brain Atlas anatomical regions coronal slice. (B) Scores from Allen Brain Atlas anatomical regions coronal slice replicate. (C) Scores from Allen Brain Atlas anatomical regions coronal slice 2, (D) Scores from Allen Brain Atlas anatomical regions sagittal anterior slice. (E) Scores from Allen Brain Atlas anatomical regions sagittal anterior slice replicate. (F) Scores from Allen Brain Atlas anatomical regions sagittal posterior slice. (G) Scores from Allen Brain Atlas anatomical regions sagittal posterior slice replicate

**Figure S7. MILWRM domain score profiles for all mouse brain samples.** (A) MILWRM-derived gene scores for MILWRM domains coronal slice. (B) Scores for MILWRM domains coronal slice replicate. (C) Scores for MILWRM domains coronal slice 2 (D) Scores for MILWRM domains sagittal anterior slice. (E) Scores for MILWRM domains sagittal anterior slice replicate.

(F) Scores for MILWRM domains sagittal posterior slice. (G) Scores for MILWRM domains sagittal posterior slice replicate
